# Casein kinase 1 controls components of a TORC2 signaling network in budding yeast

**DOI:** 10.1101/2024.01.30.578072

**Authors:** Rafael Lucena, Akshi Jasani, Steph Anastasia, Douglas Kellogg, Maria Alcaide-Gavilan

## Abstract

Tor kinases play diverse and essential roles in control of nutrient signaling and cell growth. Tor kinases are assembled into two large multiprotein complexes referred to as Tor Complex 1 and Tor Complex 2 (TORC1 and TORC2). In budding yeast, TORC2 controls a signaling network that relays signals regarding carbon source that strongly influence growth rate and cell size. However, the mechanisms that control TORC2 signaling are poorly understood. Activation of TORC2 requires Mss4, a phosphoinositol kinase that initiates assembly of a multi-protein complex at the plasma membrane that recruits and activates downstream targets of TORC2. Localization of Mss4 to the plasma membrane is controlled by phosphorylation and previous work suggested that yeast homologs of casein kinase 1γ, referred to as Yck1 and Yck2, control phosphorylation of Mss4. Here, we generated a new analog-sensitive allele of *YCK2* and used it to test whether Yck1/2 influence signaling in the TORC2 network. We found that multiple components of the TORC2 network are strongly influenced by Yck1/2 signaling.

## Introduction

Cell growth is a highly regulated process that requires the orchestration of myriad pathways that are critical for the survival and function of eukaryotic cells. The rate of growth must be tightly linked to nutrient availability to ensure that growth rate is matched to the availability of energy and biosynthetic precursors. Nutrient availability also influences the extent of growth. Thus, the amount of growth required for cell cycle progression is reduced in poor nutrients, which leads to a large reduction in cell size (Fantes and Nurse, 1977; Ferrezuelo et al., 2012; Johnston et al., 1977; Leitao and Kellogg, 2017; Schaechter et al., 1958).

Tor kinases play central roles in the mechanisms that control cell growth. Tor kinases are assembled into two large multiprotein complexes referred to as Tor Complex 1 and Tor Complex 2 (TORC1 and TORC2) (Barbet et al., 1996; Heitman et al., 1991; Loewith et al., 2002). TORC1 has been extensively studied and relays nutrient-dependent signals that control metabolism, ribosome biogenesis, autophagy and cell cycle progression (De Virgilio and Loewith, 2006; Moreno-Torres et al., 2015; Wullschleger et al., 2006). Much less is known about TORC2 (Riggi et al., 2020). In budding yeast, a complex signaling network that surrounds TORC2 is required for normal control of cell growth and size. Loss of function of TORC2 network components leads to cell size defects, as well as a failure to modulate growth rate and cell size in response to changes in nutrient availability (Lucena et al., 2018).

A key downstream target of TORC2 signaling is a pair of redundant protein kinase paralogs referred to as Ypk1 and Ypk2, which are the yeast homologs of mammalian SGK kinases (Casamayor et al., 1999; Kamada et al., 2005). To be activated by TORC2, Ypk1/2 are recruited to the plasma membrane via assembly of a multi- protein complex. A phosphoinositol kinase referred to as Mss4 converts PI_4_P to PI_4,5_P, which recruits Ypk1/2 to the TORC2 complex where they are directly phosphorylated by TORC2 (Audhya et al., 2004; Berchtold and Walther, 2009; Niles et al., 2012).

TORC2 signaling is strongly modulated by nutrient availability. For example, TORC2 signaling to Ypk1/2 is high when cells are growing in a rich carbon source like dextrose, and much lower in low quality carbon sources like glycerol and ethanol (Lucena et al., 2018). The mechanisms by which TORC2 signaling is modulated by carbon source are unknown. A potential clue came from our finding that Mss4 undergoes multi-site phosphorylation that is strongly regulated by carbon source. Thus, Mss4 is hyperphosphorylated in rich carbon and hypophosphorylated in poor carbon (Lucena et al., 2018). Furthermore, increased phosphorylation of Mss4 appears to drive recruitment of the Mss4 to the plasma membrane where it can activate TORC2. Together, these observations suggest a model in which phosphorylation of Mss4 is an important mechanism by which nutrient-availability modulates TORC2 activity (Audhya and Emr, 2003; Lucena et al., 2018). Testing this model will require a better understanding of the signals that control Mss4 phosphorylation, which are largely unknown.

Here, we have investigated the mechanisms that control phosphorylation of Mss4. A previous study suggested that Mss4 could be phosphorylated by members of the casein kinase 1γ family (CK1γ), which are encoded in yeast by a pair of redundant kinase paralogs referred to as Yck1 and Yck2 (Audhya and Emr, 2003). Cells that lack either paralog are viable, but loss of both is lethal. Yck1/2 are thought to influence diverse cellular processes, such as cell growth, endocytosis, vesicle trafficking, cell cycle progression and glucose sensing (Gadura et al., 2006; Hassaninasab et al., 2019; Moriya and Johnston, 2004; Panek, 1997; Pasula et al., 2010; Robinson et al., 1993; Snowdon and Johnston, 2016; Stalder et al., 2016).

To further investigate the functions of Yck1/2, we generated an analog-sensitive allele of YCK2 (*yck2-as1*) that can be rapidly and specifically inhibited in vivo. We used this new tool to test whether Yck1/2 influence phosphorylation of Mss4 and TORC2 signaling.

## Results

### Generation of an analog-sensitive allele of casein kinase 1

To explore the functions of Yck1/2, we generated an analog-sensitive allele of *YCK2* (*yck2-as1*) that can be rapidly and specifically inhibited *in vivo*. Cells carrying the *yck2-as1* allele in a *yck1Δ* background grew slightly slower than wild type cells at 25°C and 30°C, and substantially slower at 37°C, which indicates that the altered ATP-binding pocket causes reduced activity, as seen for other analog-sensitive alleles (Figure 1A). The *yck2-as1 yck1Δ* cells showed strong sensitivity to low nanomolar concentrations of 3-MOB-PP1, 3-BrB-PP1 and 3-MB-PP1 (Figure 1B). For further experiments, we used 3-MOB-PP1. Addition of 3-MOB-PP1 to *yck2-as1 yck1Δ* cells for 4 hours caused cells to become larger and buds became elongated (Figure 1C). Similar defects were previously observed in *yck2-ts yck1Δ* cells grown at the restrictive temperature (Robinson et al., 1993).

**Figure 1:**
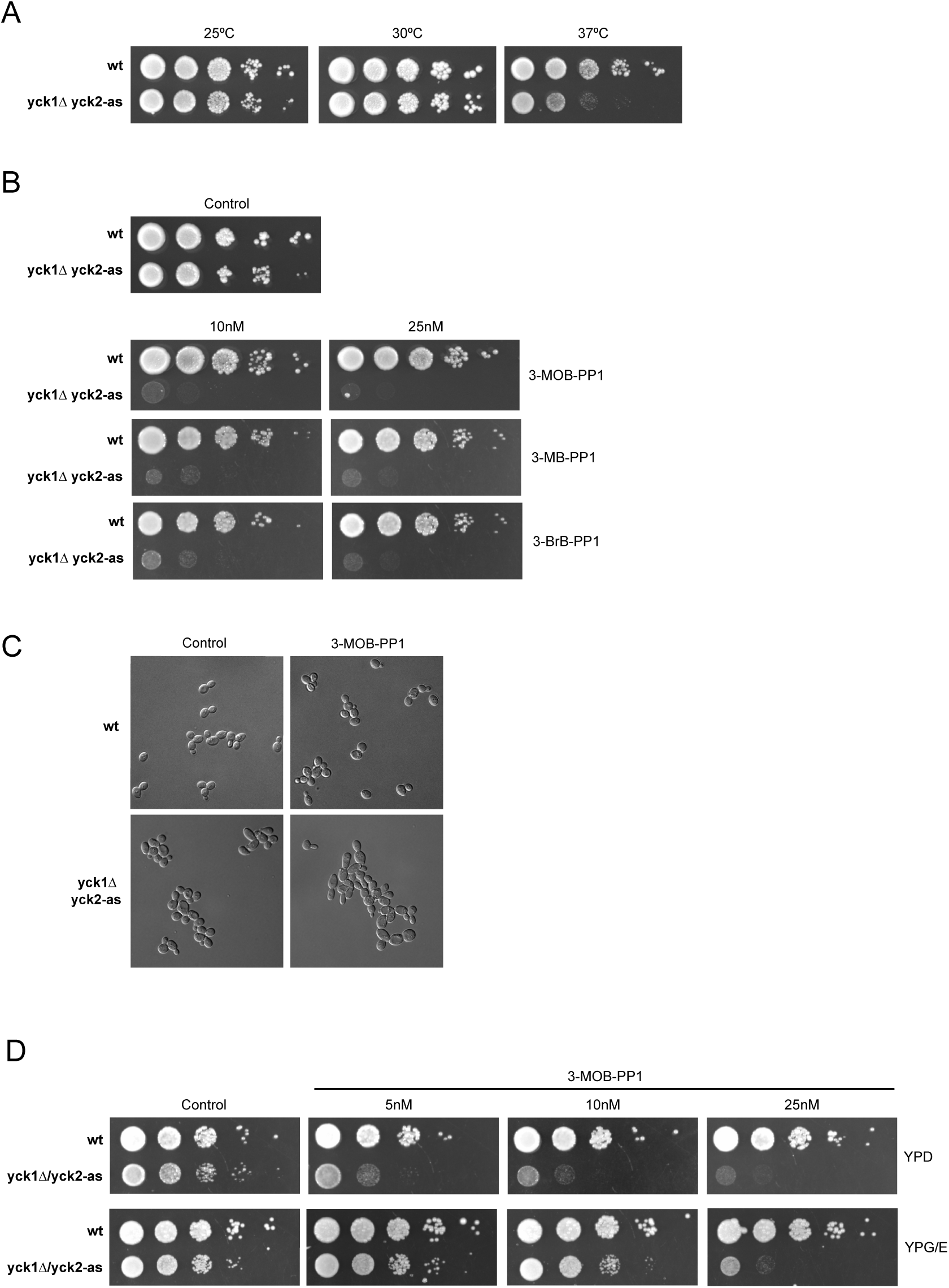
Characterization of an analog-sensitive allele of *YCK2*. (A) A series of ten-fold dilutions of wild-type and *yck1Δ yck2-as1* cells were grown in YPD medium at 25°C, 30°C and 37°C. (B) Series of ten-fold dilutions of the indicated strains were grown at 25°C in YPD medium in the absence (control) or presence of different concentrations (10 nM or 25 nM) of the indicated inhibitors (3-MOB-PP1, 3- MB-PP1 and 3-BrB-PP1) (C) Wildtype and *yck1Δ yck2-as1* cells were grown to log phase in YPD liquid medium at 25°C in the absence or presence of 0.5 µM of the 3-MOB-PP1 inhibitor, and images were taken using a Leica DM8000B microscope. (D) Series of ten-fold dilutions of wildtype and *yck1Δ yck2-as1* cells were grown at 25°C in the absence or presence of different concentrations of 3-MOB-PP1 either in YPD or YEP/GE.

Previous work suggested that Yck1/2 play roles in glucose signaling (Moriya and Johnston, 2004). Here, we found that *yck2-as1 yck1Δ* cells were more resistant to 3- MOB-PP1 in poor carbon medium (YP+2% glycerol and 2% ethanol) compared to the same concentration of analog inhibitor in rich carbon medium, consistent with an increased requirement for Yck1/2 activity in glucose (Figure 1D).

### Casein kinase I and PP2A^Rts1^ play opposing roles in regulation of Mss4

Previous work suggested that Yck1/2 regulate Mss4 phosphorylation (Audhya and Emr, 2003). These studies utilized a temperature sensitive allele of *YCK2* in a *yck1Δ* background and found that inactivation of *yck2-ts* for 30 minutes at the restrictive temperature caused loss of Mss4 phosphorylation, as measured via incorporation of ^32^P into Mss4 in vivo. Previous work also suggested that hyperphosphorylation of Mss4 is restrained by PP2A^Rts1^ (Lucena et al., 2018). We therefore used the *yck2-as1* allele to evaluate the roles of Yck1/2 and PP2A^Rts1^ in regulation of Mss4. We used western blotting to detect changes in Mss4 electrophoretic mobility caused by phosphorylation, which provides greater resolution than ^32^P incorporation.

We compared phosphorylation of Mss4 in wild type, *rts1Δ*, *yck2-as1 yck1Δ*, and *yck2-as1 yck1Δ rts1Δ* cells before and after addition of analog inhibitor. Mss4 was hyperphosphorylated in *rts1Δ* cells, as previously reported (Figure 2A, compare lanes 1 and 2). Mss4 phosphorylation was strongly reduced in *yck2-as1 yck1Δ* cells in the absence of inhibitor, consistent with reduced kinase activity of the mutant (Figure 2A, compare lanes 1 and 3). Addition of 3-MOB-PP1 caused further loss of the phosphorylated forms of Mss4 (Figure 2A, compare lanes 3 and 4). *rts1Δ* caused an increase in Mss4 phosphorylation in the *yck2-as1 yck1Δ* cells in the absence of inhibitor (Figure 2A, compare lanes 3 and 7). Addition of 3-MOB-PP1 caused rapid loss of Mss4 phosphorylation in the *yck2-as1 yck1Δ rts1Δ* cells.

**Figure 2:**
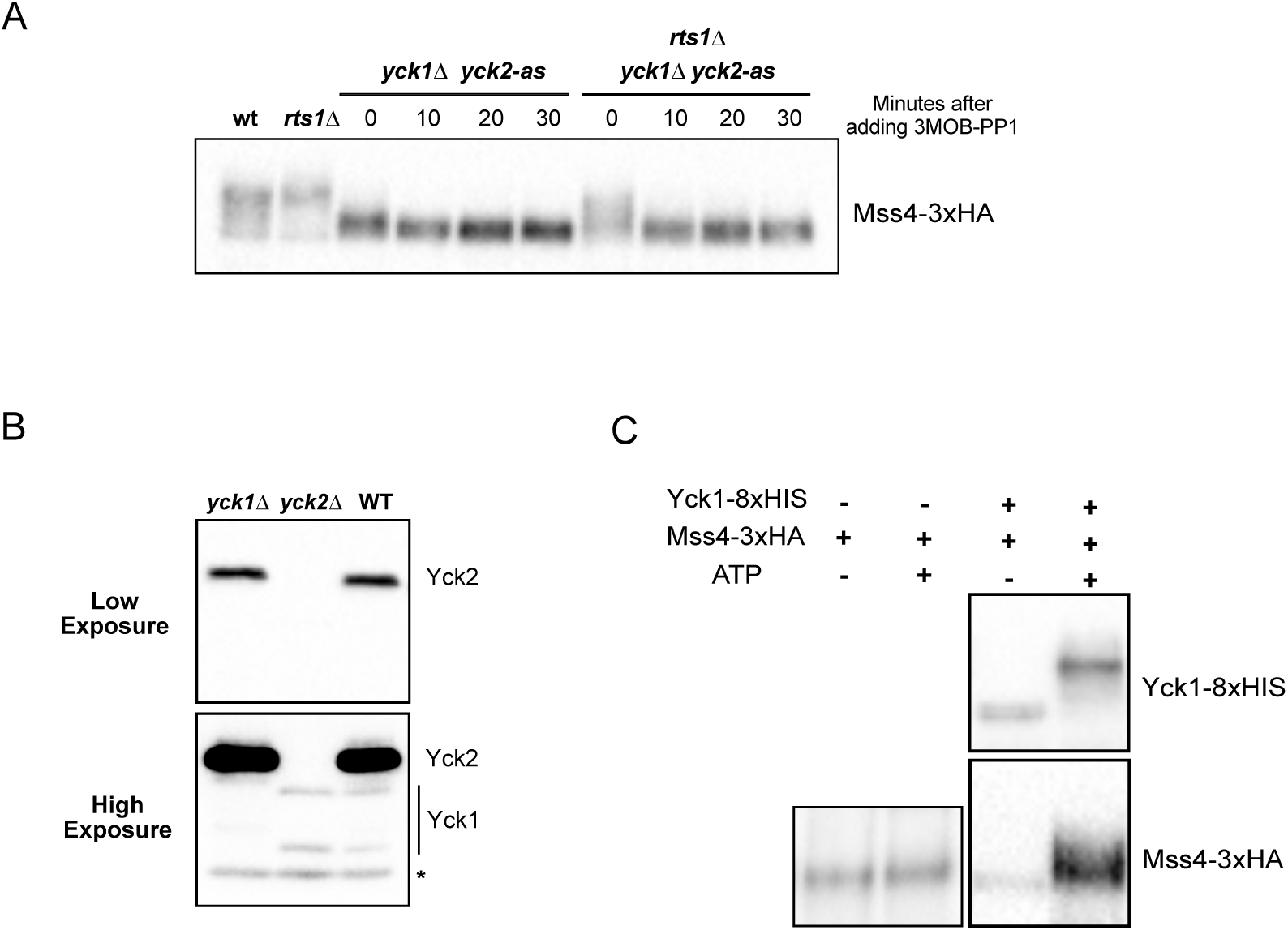
Yck1/2 are required for normal phosphorylation of Mss4. (A) Cells of the indicated genotypes were grown in YPD medium to early log phase at 25°C and 0.5 µM of 3-MOB-PP1 was then added to all *yck2-as1* cells and samples were collected at the indicated time points. Mss4-3xHA was detected by western blot. (B) Extracts were made from strains of the indicated genotypes and used for western blotting using a polyclonal anti-Yck2 antibody. Two different exposures are shown. Background bands are marked with an asterisk. (C) Kinase reactions containing affinity purified Yck1-8xHIS and Mss4-3xHA in the indicated combinations were initiated by addition of ATP and performed for 30 minutes. Ypk1 and Mss4 phosphorylation was assayed by western blot.

These data confirm that Yck1/2 strongly influence Mss4 phosphorylation. To investigate further, we raised an antibody that recognizes Yck2. We first compared the pattern of bands in wild type cells to those seen in *yck2Δ* and *yck1Δ* cells to define which bands correspond to Yck2, which showed that the antibody strongly detects Yck2 and weakly detects Yck1 (Figure 2B). We found that a Yck1-8xHIS fusion protein purified from insect cells was able to induce partial hyperphosphorylation of Mss4 in vitro, consistent with the possibility that Yck1/2 directly phosphorylates Mss4 (Figure 2C). However, the data do not rule out models in which Yck1/2 activate another kinase that phosphorylates Mss4 or inhibit a phosphatase that acts on Mss4.

The fact that loss of Rts1 does not cause quantitative hyperphosphorylation of Mss4 and that Mss4 dephosphorylation occurs when Yck1/2 is inhibited in *rts1Δ* cells, suggests that multiple phosphatases work on Mss4.

### PP2A influences Yck1/2 phosphorylation *in vivo* and *in vitro*

We found that PP2A^Rts1^ purified from yeast was not able to dephosphorylate Mss4 *in vitro*, even though it was active against a number of other substrates (not shown), which suggests that PP2A^Rts1^ acts indirectly to influence Mss4 phosphorylation. An alternative model is that PP2A^Rts1^ leads to an increase in the activity of Yck1/2. Yck2 undergoes autophosphorylation, which could promote Yck2 activity (Stalder et al., 2016). In this case, if PP2A^Rts1^ opposes autophosphorylation it would be an inhibitor of Yck1/2.

We used Yck1-8xHIS purified from insect cells and 3xHA-Yck2 purified from yeast for in vitro assays. Both underwent extensive autophosphorylation in vitro that could be detected as an electrophoretic mobility shift (Figure 3A). Furthermore, purified PP2A^Rts1^ was able to oppose autophosphorylation of Yck1 and Yck2 *in vitro* (Figure 3A).

**Figure 3:**
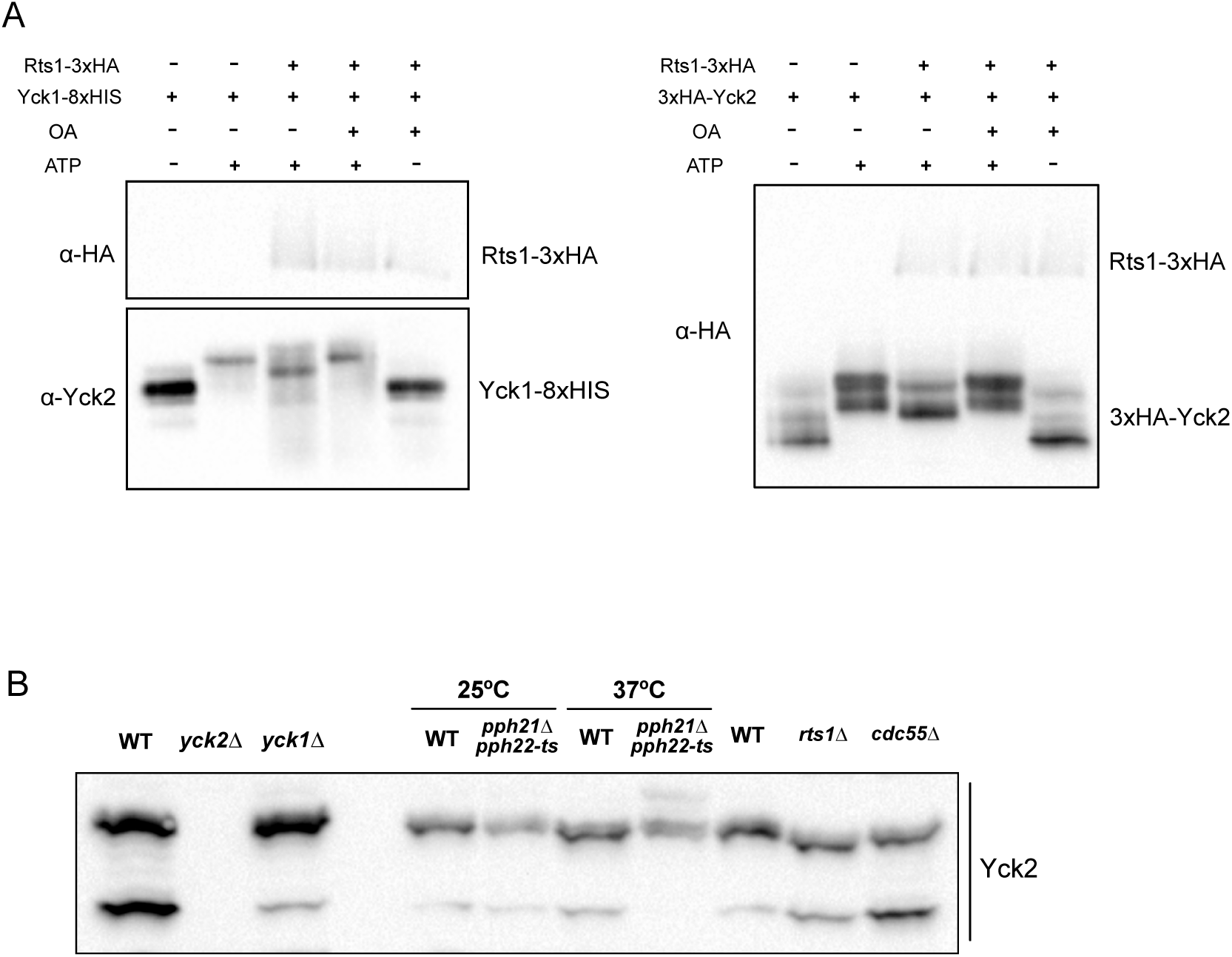
PP2A influences Yck1/2 phosphorylation. (A) *In vitro* assays containing affinity purified 3xHA-Ypk1 and either 3xHA-Yck2 or Yck1- 8xHIS in the indicated combinations were performed for 20 minutes in the presence or absence of ATP and okadaic acid (OA). Ypk1 and Yck1/2 phosphorylation was assayed by western blot. (B) Cells of the indicated genotypes were grown to log phase at 25°C in YPD medium. Wild-type and *pph21Δ pph22-ts* cells were then shifted to 37°C for 30 minutes before collecting samples. Phosphorylation of Yck2 was assayed by Phos-tag western blot.

To test whether PP2A^Rts1^ influences the phosphorylation state of Yck2 *in vivo*, we tested whether inactivation of PP2A catalytic subunits or Rts1 causes hyperphosphorylation of Yck2 in vivo. In budding yeast, PP2A is encoded by two redundant paralogs that are referred to as *PPH21* and *PPH22*. Cells that lack either paralog are viable, whereas loss of both is nearly lethal. To inactivate the catalytic subunits, we used a temperature sensitive allele of *PPH21* in a *pph22Δ* background (*pph21-172 pph22Δ,* (Evans and Stark, 1997). To obtain better resolution of phosphorylated forms of Yck2, we utilized Phos-Tag gels. We found that Yck2 in log phase wild type cells exists in two phosphorylation forms (Figure 3B). Inactivation of *PPH21* and *PPH22* caused the appearance of a hyperphosphorylated form of Yck2. However, Yck2 did not undergo hyperphosphorylation in *rts1Δ* cells (Figure 3B). Loss of Cdc55, the other B-type regulatory subunit for PP2A, also had no effect on Yck2 phosphorylation. These observations suggest that forms of PP2A that include Cdc55 or Rts1 can not be solely responsible for Yck2 dephosphorylation; however, it is possible that PP2A^Rts1^ and PP2A^Cdc55^ work redundantly to control PP2A-dependent phosphorylation of Yck2. It is also possible that another form of PP2A is responsible for directly controlling Yck2 phosphorylation. For example, previous studies have shown that PP2A catalytic subunits can also associate with *TAP42*, which is thought to be regulated by nutrient-dependent signals (Di Como and Arndt, 1996).

### Yck1/2 influence TORC2 signaling

Since Yck1/2 are required for normal phosphorylation of Mss4, we next tested whether they influence signaling events in the TORC2 signaling network. Correlative data suggest that phosphorylation of Mss4 drives increased localization of Mss4 to the plasma membrane, as well as increased TORC2-dependent activation of Ypk1/2. Thus, loss of Mss4 phosphorylation caused by *yck2-as1 yck1Δ* would be expected to cause decreased TORC2-dependent phosphorylation of Ypk1/2.

To analyze the effects of inhibiting Yck1/2 activity on TORC2 signaling, we utilized a phosphospecific antibody that detects a TORC2-dependent phosphorylation site that is present on Ypk1 (T662P, (Niles et al., 2012). The site is also present on Ypk2 so the antibody recognizes both proteins. In addition to TORC2-dependent phosphorylation at T662, Ypk1 undergoes multi-site phosphorylation by multiple kinases that cause electrophoretic mobility shifts. Therefore, we also used an antibody that recognizes Ypk1, which allowed us to detect phosphorylation of Ypk1 at sites other than T662 that cause changes in electrophoretic mobility. We found that T662 phosphorylation was modestly reduced in *yck2-as1 yck1Δ* cells in the absence of inhibitor (Figure 4A). Addition of 3-MOB-PP1 did not cause a substantial further reduction in T662P levels (Figure 4A). This result is similar to the results obtained for Mss4, where *yck2-as1* caused a loss of phosphorylation of Mss4 in the absence of inhibitor and addition of inhibitor caused only minimal further effects. Analysis of Ypk1 phosphorylation with the anti-Ypk1 antibody showed that *yck2-as1* in the absence of inhibitor caused a reduction in phosphorylation of Ypk1. Prolonged incubation in the presence of the inhibitor caused further changes in the phosphorylation state of Ypk1 (Figure 4A).

**Figure 4:**
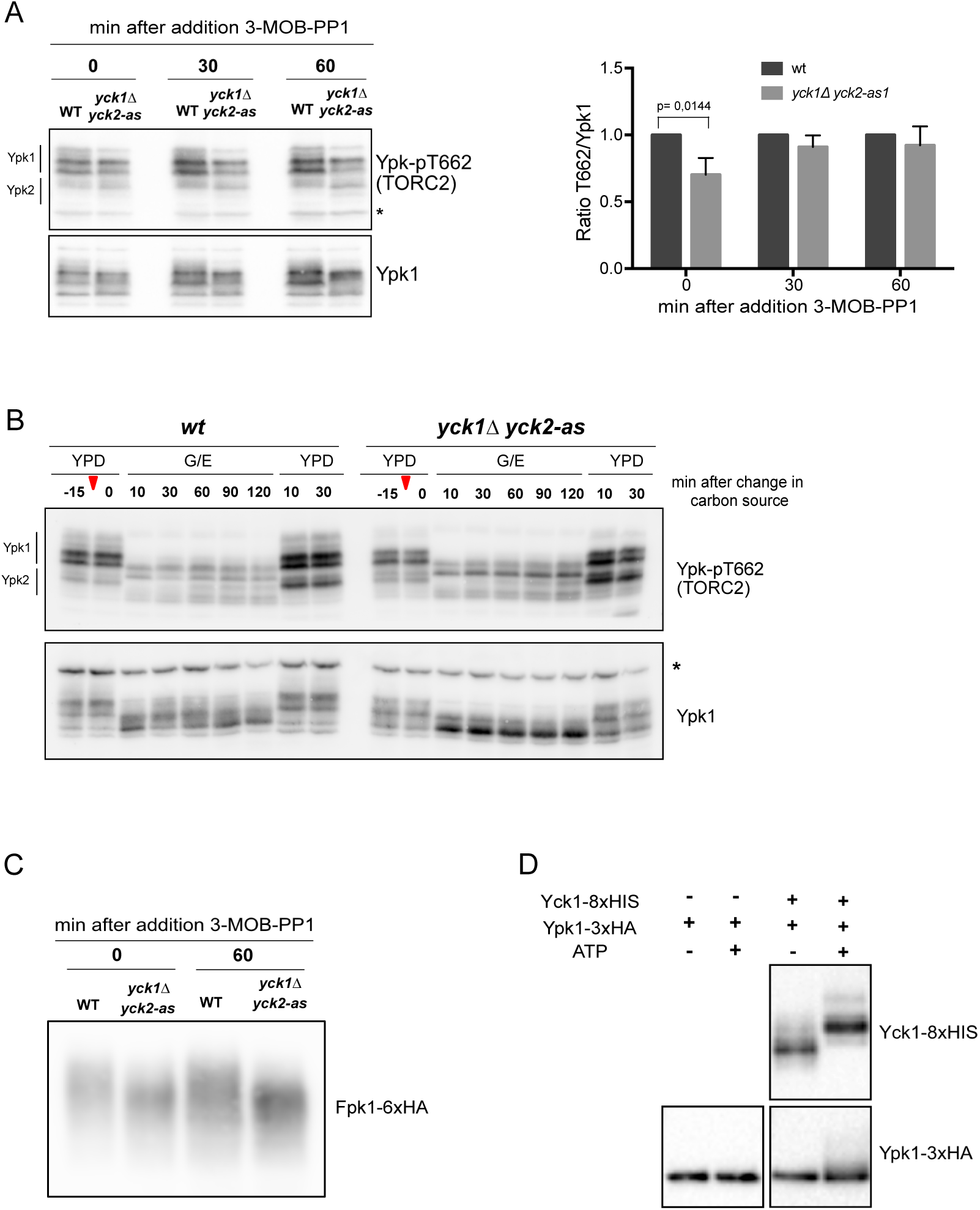
Casein kinase 1 regulates TORC2 signaling pathway. (A) Wildtype and *yck1Δ yck2-as1* were grown to early log phase at 25°C and 0.5 µM of 3-MOB-PP1 was the added to both strains and samples were collected at the indicated time points. Ypk-pT662 and Ypk1 were analyzed by western blot. Quantification of the Ypk-pT662 signal over total Ypk1 protein is shown. Error bars represent the SD of the mean of three biological replicates. (B) Wildtype and *yck1Δ yck2-as1* cells were grown to early log phase at 25°C and 0.5 µM of 3-MOB-PP1 was then added to both strains for 15 minutes and cultures were then shifted from YPD to YPG/E medium containing the same amount of inhibitor. After 2 hours in YPG/E medium cells were shifted back to YPD medium (indicated by red arrows). Samples were collected at the indicated time points and Ypk-pT662 and Ypk1 were analyzed by Western blotting. (C) Wildtype and *yck1Δ yck2-as1* were grown to early log phase at 25°C and 0.5 µM of 3-MOB-PP1 was then added to both strains and samples were collected at the indicated time points. Fpk1 phosphorylation was assayed by western blot. (D) Kinase reactions containing affinity purified Yck1-8xHIS and Gal-3xHA-Ypk1 in the indicated combinations were initiated by addition of ATP and performed for 30 minutes. Ypk1 and Yck1 phosphorylation was assayed by western blot.

In previous work we found that a shift from rich to poor carbon causes a rapid and dramatic decrease in TORC2-dependent phosphorylation of Ypk1/2, as well a dramatic decrease in overall Ypk1 phosphorylation (Lucena et al., 2018). Therefore, we next tested whether Yck1/2 are required for modulation of TORC2 signaling in response to changes in carbon source. We again saw that *yck2-as1 yck1Δ* cells showed reduced T662 phosphorylation in rich carbon medium before the shift. The response of T662 phosphorylation to a shift to poor nutrients appeared to occur normally (Figure 4B). However, inhibition of *yck2-as1* caused major changes in the electrophoretic mobility of Ypk1 that indicated a loss of phosphorylation at sites other than T662 when cells were shifted to poor carbon.

Since Yck1/2 are thought to be involved in glucose signaling we also tested whether TORC2 signaling and Ypk1 respond normally when cells are shifted from poor carbon back to glucose (Figure 4B, last two lanes). The response appeared to be normal, again indicating that Ypk1/2 are not required for the response of TORC2 signaling to changes in carbon source.

Phosphorylation of Ypk1 that can be detected via electrophoretic mobility shifts is due at least partly to a redundant pair of kinase paralogs called Fpk1 and Fpk2, which play poorly understood roles in TORC2 signaling (Roelants et al., 2010). To test whether Yck1/2 control Ypk1 phosphorylation via Fpk1/2, we tested whether phosphorylation of Fpk1 is dependent upon Yck1/2. We found that Fpk1 phosphorylation that can be detected via electrophoretic mobility shifts was reduced in *yck2-as1 yck1Δ* cells, consistent with the possibility that Yck1/2 influence Ypk1 phosphorylation via Fpk1/2 (Figure 4C).

Another possibility is that Yck1/2 directly influence Ypk1 phosphorylation. We found that purified Yck1-8xHIS appeared to be able to phosphorylate purified Ypk1, as evidenced by a slight shift in electrophoretic mobility (Figure 4D). An ability of Yck1/2 to regulate Ypk1/2 could explain effects on Fpk1/2, since it has been proposed that Ypk1/2 regulate Fpk1/2 (Roelants et al., 2010).

### Yck1/2 are required for normal regulation of Rts1 phosphorylation and show genetic interactions with Rts1

The TORC2 signaling network influences phosphorylation of Rts1 via poorly understood feedback signals. For example, a shift from rich to poor carbon causes a dramatic hyperphosphorylation of Rts1 that is dependent upon Ypk1/2, yet Rts1 is also required for normal regulation of Ypk1/2 (Alcaide-Gavilán et al., 2018; Lucena et al., 2018). Although the functions of Rts1 phosphorylation are largely unknown, it provides a readout of signaling events associated with the TORC2 signaling network. To test whether Yck1/2 influence Rts1 phosphorylation, we shifted wild type control cells and *yck2-as1 yck1Δ* cells from rich to poor carbon and inhibited *yck2-as1* with 3-MOB-PP1 at the time of the shift (Figure 5A). Previous work has shown that all of the electrophoretic mobility shifts of Rts1 that can be detected by SDS -PAGE are due to phosphorylation (Alcaide-Gavilán et al., 2018). Rts1 phosphorylation was already reduced before the shift to poor carbon and before addition of inhibitor (Figure 5A, compare the zero timepoints). Furthermore, hyperphosphorylation of Rts1 failed to occur when the *yck2-as1 yck1Δ* cells were shifted to poor carbon. Inactivation of Ypk1/2 has similar effects on hyperphosphorylation of Rts1 in response to a shift from rich to poor carbon, consistent with a model in which Yck1/2 influence Ypk1/2 activity via Mss4 (Alcaide-Gavilán et al., 2018).

**Figure 5:**
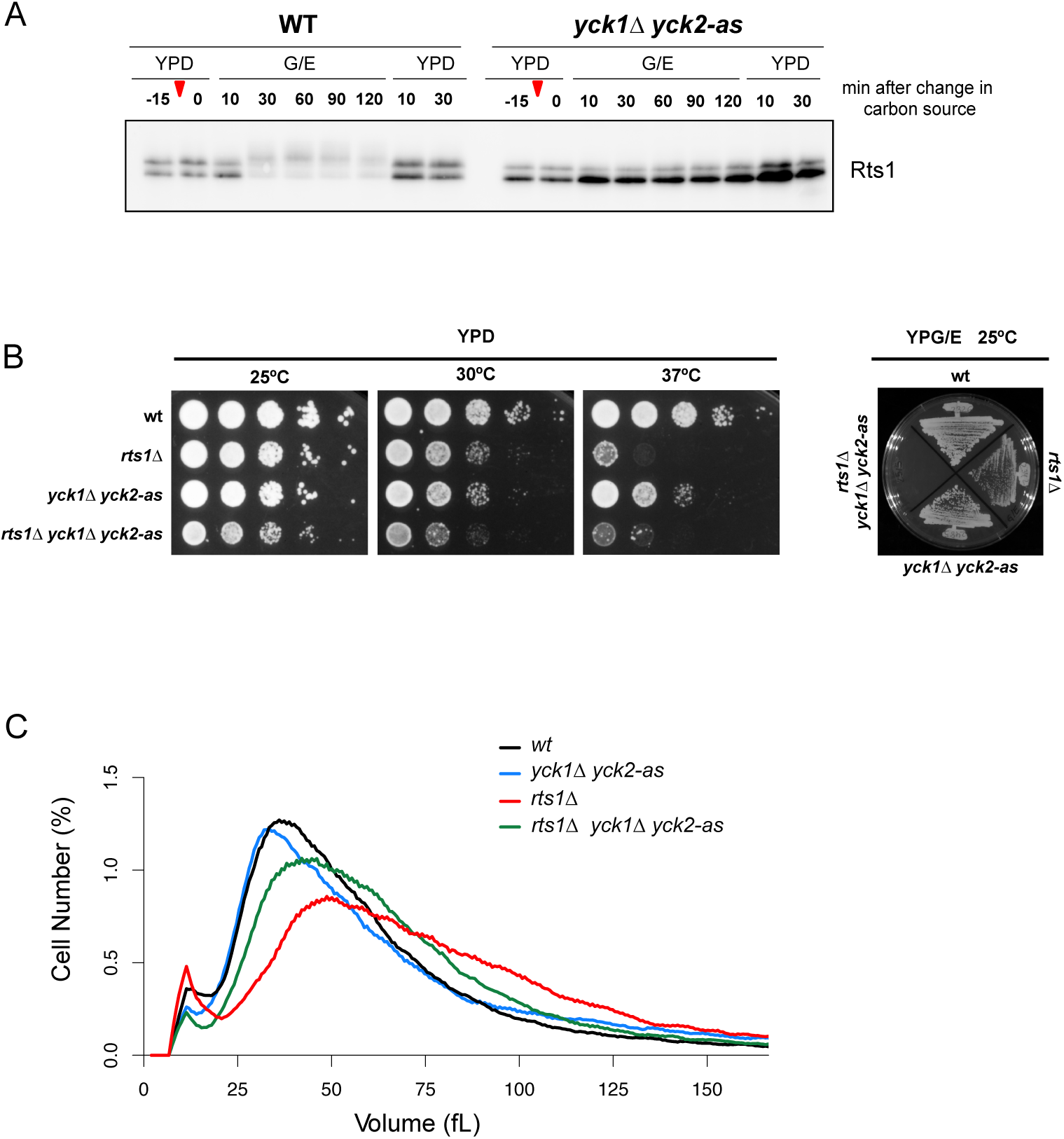
Yck1/2 and Rts1 show genetic interactions. (A) Wildtype and *yck1Δ yck2-as* cells were grown to early log phase at 25°C and 0.5uM of 3-MOB-PP1 was then added to both strains for 15 minutes and cultures were shifted from YPD to YPG/E medium containing the same amount of inhibitor (indicated by red arrows). After 2 hours in YPG/E medium cells were shifted back to YPD medium. Samples were collected at the indicated time points and Rts1 was analyze by western blot. (B) Series of 10-fold dilutions of the indicated strains were grown at different temperatures in YPD medium. Cells of the indicated genotypes were grown in YPG/E medium at 25°C. (C) Cells of the indicated genotypes were grown to log phase at 25°C and cell-size distributions were determined using a Coulter Counter. Each plot is the average of three biological replicates. For each biological replicate, three technical replicates were analyzed and averaged.

**Figure 6:**
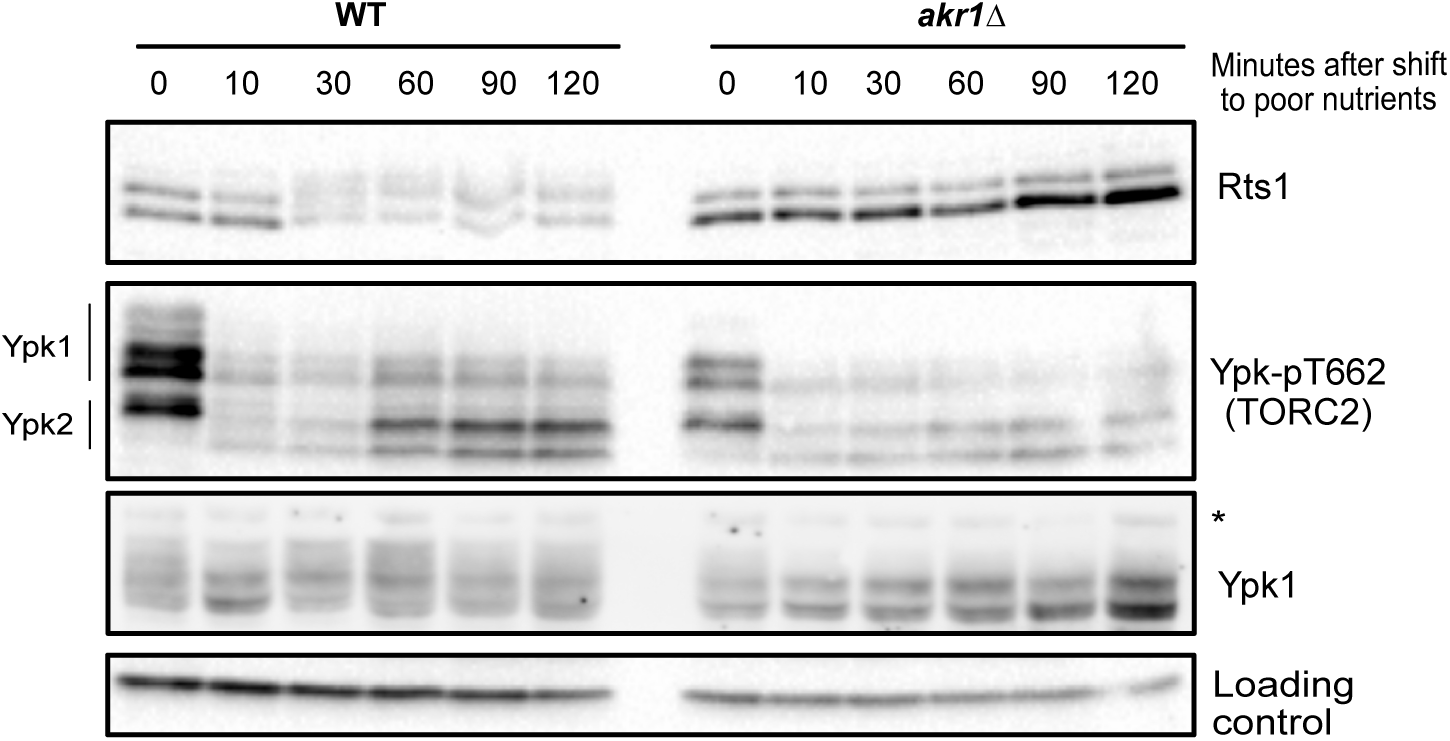
Membrane localization of Casein kinase 1 is important for normal regulation of the TORC2 signaling pathway. Wild-type and *akr1Δ* cells were grown to early log phase at 25°C in YPD medium. Cells were washed into YPG/E medium and samples were collected at the indicated time points. The behavior of Ypk-pT662 and Ypk1 was assayed by western blot at the indicated times.

We next tested for genetic interactions between Yck1/2 and PP2A^Rts1^. We found that *yck2-as1 yck1Δ* slightly reduced the rate of proliferation of *rts1Δ* cells in the absence of inhibitor (Figure 5B). Furthermore, *yck2-as1 yck1Δ* cells caused a partial rescue of *rts1Δ* cell size defects in rich media, but a synthetic lethal effect in poor media (Figure 5B, 5C and D). These genetic interactions are consistent with the idea that Yck1/2 and Rts1 share related functions.

### Membrane localization is required for the ability of Yck1/2 to influence components of the TORC2 network

Yck1/2 are anchored in the plasma membrane via a palmitoyl group that is added by a palmitoyl transferase called Akr1 (Babu et al., 2004; Feng and Davis, 2000; Pasula et al., 2010). Cells that lack Akr1 are viable but show defects in cell size and shape that are similar to the effects of inactivating Yck1/2, which indicates that some functions of Yck1/2 are dependent upon membrane localization. The fact that loss Yck1/2 is lethal, whereas loss of Akr1 is not, indicates that not all functions of Yck1/2 are dependent upon membrane localization. We found that *akr1Δ* caused decreased Ypk1-T662 phosphorylation in rich carbon but did not cause a failure in the response of Ypk1-T662 phosphorylation to a shift from rich to poor carbon, as we found for inhibition of *yck2- as1*. In addition, *akr1Δ* caused defects in hyperphosphorylation of Rts1 in response to a shift to poor carbon that were similar to the effects of *yck2-as1*. These results suggest that membrane localization of Yck1/2 is required for their effects on the TORC2 network. However, since Akr1 also palmitoylates several other proteins, it is possible that loss of membrane localization of several proteins is responsible for the effects of *akr1Δ*.

## DISCUSSION

We developed an analog-sensitive allele of *YCK2* that provides a new tool for analysis of the functions of Yck1/2 in yeast. As a first application, we used the *yck2-as1* allele to analyze how Yck1/2 and PP2A^Rts1^ influence phosphorylation of Mss4. We found that loss of Yck1/2 activity causes a large decrease in phosphorylation of Mss4 that can be detected by electrophoretic mobility shifts. Since Mss4 and Yck1/2 are both localized to the plasma membrane, it is possible that Yck1/2 directly phosphorylate Mss4. We found that purified Yck1 was able to phosphorylate Mss4 in vitro, consistent with the possibility that it directly phosphorylates Mss4 in vivo.

Loss of Rts1 causes hyperphosphorylation of Mss4 but does not lead to quantitative full hyperphosphorylation of Mss4, which indicates that PP2A^Rts1^ can not be the only phosphatase that works on Mss4. We found that loss of Rts1 causes a partial recovery of Mss4 phosphorylation in *yck2-as1 yck1Δ* cells. There are two models that could explain the data. In one model, there are multiple phosphatases that work on the same sites on Mss4. In this case, loss of Rts1 would remove one of the phosphatases so that the other phosphatases do not provide sufficient activity to fully dephosphorylate Mss4. An alternative model is that there are multiple kinases and phosphatases that each work on distinct sets of sites on Mss4. In this case, PP2A^Rts1^ could oppose a kinase that is not Yck1/2, and loss of Rts1 in *yck2-as1 yck1Δ* cells would still lead to hyperphosphorylation of Mss4. It is important to note that detection of phosphorylation events via electrophoretic mobility shifts in standard SDS-PAGE provides limited resolution, as some phosphorylation events cause no shift and others can cause large or small shifts alone or in combination with other events.

PP2A^Rts1^ purified from yeast was not able to dephosphorylate Mss4 *in vitro*, which suggests that it acts indirectly. As an alternative we considered a model in which loss of PP2A^Rts1^ leads to an increase in the activity of Yck1/2. We tested this idea and found that PP2A^Rts1^promotes dephosphorylation of Yck1 and Yck2 *in vitro*. However, only the inactivation of PP2A catalytic subunit causes hyperphosphorylation of Yck2 *in vivo*. This result suggest PP2A^Rts1^ can not be solely responsible for Yck2 dephosphorylation. It is possible that both PP2A regulatory subunits (Cdc55 and Rts1) work redundantly on Yck2. Alternatively, another form of PP2A could be responsible for directly controlling Yck2 phosphorylation.

Phosphorylation of Mss4 is thought to help recruit it to the plasma membrane to promote TORC2 signaling (Audhya et al., 2004; Berchtold and Walther, 2009; Niles et al., 2012). In support of this, we previously found that loss of Rts1 leads to hyperphosphorylation of Mss4, increased recruitment of Mss4 to the plasma membrane, and increased TORC2-dependent phosphorylation of Ypk1/2 (Lucena et al., 2018). It was therefore surprising to find that loss of Yck1/2 activity caused relatively modest effects on TORC2-dependent phosphorylation of Ypk1/2. A potential explanation is that a kinase other than Yck1/2 is responsible for phosphorylation-dependent localization of Mss4. Alternatively, *yck2-as1* could retain sufficient activity in the presence of inhibitor to permit a low level of Mss4 phosphorylation that is sufficient for localization.

Previous work suggested that Yck1/2 play roles in a glucose sensing pathway (Gadura et al., 2006; Kim et al., 2022; Pasula et al., 2010; Robinson et al., 1992; Snowdon and Johnston, 2016). For example, there is evidence that Yck1/2 are required for normal membrane localization and stability of the Rgt2 glucose sensor, or that Yck1/2 relay a glucose signal to Rgt2. Here, we found that loss of Yck1/2 activity had no effect on the response of TORC2 signaling to changes in carbon source. Thus, in *yck2-as1 yck1Δ* cells TORC2-dependent phosphorylation of T662 of Ypk1 responded normally to a shift from glucose to poor carbon, and to a shift from poor carbon to glucose, which indicates that glucose signaling to TORC2 is not dependent upon normal activity of Yck1/2. However, we found that there is a greater requirement for Yck1/2 activity when cells are growing in glucose containing medium compared to poor carbon medium, consistent with previous studies that found a role for Yck1/2 in pathways that mediate responses to glucose.

We also tested whether loss of Yck1/2 affects phosphorylation of Ypk1 that can be detected via electrophoretic mobility shifts. Previous studies suggested that much of the phosphorylation of Ypk1 that can be detected via electrophoretic mobility shifts is dependent upon the Fpk1/2 kinases (Roelants et al., 2010). We found that loss of Yck1/2 causes a loss of Ypk1 phosphorylation that was particularly apparent when cells were shifted from rich to poor carbon. We further discovered that loss of Yck1/2 causes a substantial loss of Fpk1 phosphorylation, which suggests that Yck1/2 could influence Ypk1 phosphorylation via activation of Fpk1/2. The functions of Fpk1/2 are poorly understood. Cells that lack Fpk1/2 are viable but have hyperpolarized actin in the growing bud, which suggests that Fpk1/2 control organization of the actin cytoskeleton, potentially via regulation of TORC2 signaling (Rispal et al., 2015). Fpk1/2 are also thought to control phospholipid flippases at the plasma membrane (Roelants et al., 2010). An alternative hypothesis is that Yck1/2 directly phosphorylate Ypk1. We tested this idea and found that a purified Yck1 was able to phosphorylate Ypk1 in vitro, suggesting that Yck1 could contribute to Ypk1 phosphorylation in vivo, and that loss of Fpk1 phosphorylation in *yck2-as1 yck1Δ* cells could be due to loss of Ypk1 activity.

Finally, we tested whether Yck1/2 activity influences phosphorylation of Rts1. In wild type cells growing in rich carbon, Rts1 appears to be equally distributed between a phosphorylated form and a dephosphorylated form, and a shift from rich to poor carbon causes further hyperphosphorylation of Rts1. In *yck2-as1 yck1Δ* cells, Rts1 exists primarily in the dephosphorylated form and completely fails to undergo hyperphosphorylation when cells are shifted to poor carbon. Moreover, we found genetic interactions between Rts1 and Yck1/2 that are consistent with the idea that they share related functions.

The data thus far can not be explained by a simple model. It is clear that inhibition of Yck1/2 results in a strong loss of phosphorylation of Mss4, Rts1, Ypk1/2 and Fpk1/2, but it does not appear that Yck1/2 can directly phosphorylate all of these proteins, and it is not yet possible to connect all of these events to a single regulatory step controlled by Yck1/2. It is possible that Yck1/2 influences the phosphorylation of each of these TORC2 components via different mechanisms. More work is therefore needed to define the mechanisms and functions of Yck1/2 signaling within the TORC2 network.

## MATERIALS AND METHODS

### Yeast strains and media

All strains are in the W303 background (leu2-3,112 ura3-1 can1-100 ade2-1 his3- 11,15 trp1-1 GAL + ssd1-d2) with the exception of strain LRB1039 which is in an unknown background. Table S1 shows additional genetic features. One-step PCR-based gene replacement was used for making deletions and adding epitope tags at the endogenous locus (Janke et al., 2004; Longtine et al., 1998). Cells were grown in YP medium (1% yeast extract, 2% peptone, 40 mg/liter adenine) supplemented with 2% dextrose (YPD), 2% galactose (YPGal), or 2% glycerol/ethanol (YPG/E). For experiments using analog-sensitive alleles, cells were grown in YPD medium without supplemental adenine. For nutrient shifts, cells were grown in YPD medium overnight to log phase. Cells were then washed three times with YPG/E medium and resuspended in YPG/E medium.

To create the *yck2-as1* allele, we first amplified the YCK2 coding sequence along with upstream and downstream non-coding regions and cloned into the BamHI and EcoRI sites of yIPlac211 (oligos: gcgggatccCCGCATATATTCCTAAGTACCTTTTTTTTCAGACAG and gcggaattcCTTGATACTCTGTATTTAGTACACAATAACGCCGACG). Site directed mutagenesis was then used to mutate L149 to G and L90 to I to create pYck2-as-GI. This plasmid was cut with MscI to target integration at Yck2 in a *yck1Δ* background. After selection of transformants, the strain was grown in YPD and plated on FOA to select for recombination events that looped out the plasmid. To identify recombination events that left the mutants in YCK2, the isolates were screened for sensitivity to 3- MOB-PP1 and sequenced to verify the presence of the mutations.

The adenine analog inhibitors 3-MOB-PP1, 3-MB-PP1 and 3-BrB-PP1 stock were prepared in 100% DMSO at 10 mM and added to cultures at a final concentration of 5, 10 and 25 nM. 3-MOB PP1, 3-MB-PP1 and 3-BrB-PP1 inhibitors were a gift from Kevan Shokat (UCSF).

### Analysis of cell size and cell proliferation assays

Cell cultures were grown overnight to early log phase at 25°C. A 900 ml sample of each culture was fixed with 100 ml of 37% formaldehyde for 30 min and then washed twice with PBS + 0.04% sodium azide + 0.02% Tween-20. Cell size was measured using a Coulter Counter Z2 (Channelizer Z2; Beckman Coulter) as previously described (Jorgensen et al 2002). In brief, cells were diluted into 10 mL diluent (Isoton II; Beckman Coulter) and sonicated for 3 s before cell sizing. Each plot is the average of three independent biological replicates in which three independent technical replicates were averaged.

To assay the rate of cell proliferation on plates, cells were grown overnight in the indicated medium at 25°C and adjusted to an OD_600_ of 0.5. Ten-fold serial dilutions were spotted onto YPD or YPG/E containing DMSO or different concentrations of 3-MOB- PP1, 3-MB-PP1 and 3-BrB-PP1 and incubated at 25°C, 30°C or 37°C.

### Microscopy

To visualize cells in figure 1C, wildtype and *yck1Δ yck2-*as1 were grown together to early log phase in the absence or presence of 0.5 µM of 3-MOB-PP1. Fields of view containing both genotypes were imaged using a Leica DM8000B microscope equipped with an objective lens (HCX PL APO 100×/1.40 OIL PH3 CS), a DFC350FX camera and LAS AF software following the instructions of the manufacturer.

### Production of polyclonal Yck2 antibody

An antibody that recognizes Yck2 was generated by immunizing rabbits with a fusion protein expressed in bacteria. A plasmid expressing full-length Yck2 was constructed with the Gateway cloning system. Briefly, a PCR product that includes the full-length open reading frame for Yck2 was cloned into the entry vector pDONR221 (pVT87). The resulting donor plasmid was used to generate a plasmid that expresses Yck2 fused at its N-terminus to 6XHis-TEV, using expression vector pDEST17 (6XHis, pVT89). The 6XHis-TEV-Yck2 fusion was expressed in BL21 cells and purified via Ni^2+^ affinity chromatography in the presence of 2M urea using standard procedures, yielding 10 mg from 4 liters of bacterial culture. A milligram of the purified protein was used to immunize a rabbit. The 6XHis-Yck2 fusion protein was coupled to Affigel 10 (Bio-Rad,Hercules,CA) to create an affinity column for purification of the antibody.

### Western blotting

To prepare samples for Western blotting, 1.6 ml of culture was collected and centrifuged at 13,000 rpm for 30 s. The supernatant was removed, and glass beads (250 µl) were added before freezing in liquid nitrogen. To analyze cells shifted from rich to poor nutrients, cultures were grown in YPD overnight at 25° to an OD_600_ of 0.8. After adjusting optical densities to normalize protein loading, cells were washed three times with a large volume of YPG/E medium and then incubated at 25° in YPG/E for the time course. Cells were lysed by bead-beating in 140 µl of 1× sample buffer (65 mM Tris-HCl, pH 6.8, 3% SDS, 10% glycerol, 50 mM NaF, 100 mM beta glycerolphosphate, 5% 2-mercaptoethanol, 2 mM phenylmethylsulfonyl fluoride [PMSF], and bromophenol blue). The PMSF was added immediately before lysis from a 100 mM stock in ethanol. Cells were lysed in a mini-beadbeater-16 (Biospec Products) at top speed for 2 min. The samples were then centrifuged for 15 s at 13,000 rpm, placed in a boiling water bath for 5 min, and centrifuged for 5 min at 13,000 rpm. SDS–PAGE was carried out as previously described (Harvey et al., 2005) at a constant current of 20 mA. Proteins were transferred to nitrocellulose using a Trans-Blot Turbo system (Bio-Rad). Blots were probed with primary antibody overnight at 4°. Proteins tagged with the HA epitope were detected with the 12CA5 anti-HA monoclonal antibody. Rabbit anti–phospho- T662 antibody was used to detect TORC2-dependent phosphorylation of YPK1/2 at a dilution of 1:10,000 in TBST (10 mM Tris-Cl, pH 7.5, 100 mM NaCl, and 0.1% Tween 20) containing 3% milk. Total Ypk1 was detected using anti-Ypk1 antibody (Alcaide-Gavilán et al., 2020) at a dilution of 1:10,000. All blots were probed with an HRP-conjugated donkey anti-rabbit secondary antibody (catalog number NA934V; GE Healthcare) or HRP-conjugated donkey anti-mouse antibody (catalog number NXA931; GE Healthcare) or HRP-conjugated donkey anti-goat antibody (catalog number sc-2020; Santa Cruz Biotechnology) for 45–90 min at room temperature. Secondary antibodies were detected via chemiluminescence with Advansta ECL reagents and a Bio-rad ChemiDoc imaging system.

Quantification of Western blot signals was performed using ImageJ (Schneider et al., 2012). Quantification of Ypk-pT662 phosphorylation was calculated as the ratio of the phospho- specific signal over the total Ypk1 protein signal, with wild-type signal normalized to a value of 1. At least three biological replicates were analyzed and averaged to obtain quantitative information.

### Immunoaffinity purifications

Immunoaffinity purification of Gal-3xHA-Ypk1 (DK2724) and Gal-3xHA-Yck2 (DK1646) were performed as follow. Cells containing a 3XHA-tagged copy of Ypk1 or Yck2 under the control of the GAL1 promoter were grown overnight at 30°C in YPG/E medium to OD_600_ = 0.5. Galactose was added to 1% and cells were incubated at 30°C for 3h. The cells were pelleted and ground under liquid nitrogen and 12 g of cell powder was resuspended in 30 ml of extract buffer containing 2 mM PMSF (50 mM Hepes-KOH, pH 7.6, 1 M KCl, 1 mM MgCl_2_, 1 mM EGTA, 5% glycerol, 0.25% Tween-20 and 2 mM PMSF). Crude extracts were stirred for 7-10 minutes at 4°C to break up cell chunks before centrifugation. All subsequent steps were performed at 4 ° C. The extract was centrifuged for 5 min at 10,000 g, followed by an additional centrifugation step at 45,000 g for 45 min. After the final spin, 15 ml of the extract was added to the anti-HA antibody beads. Beads were equilibrated by washing three times in extract buffer before the addition of extract. The clarified extracts were incubated with 650 μg of affinity purified rabbit polyclonal HA antibody bound to 500 μl protein A agarose beads for 3 hours. The beads were pelleted by centrifugation, the supernatant was removed, and the remaining extract was incubated with the beads for an additional 1.5h at 4°C. The beads were pelleted by centrifugation and washed three times with 15 ml of ice-cold extract buffer without PMSF. After the final wash, the beads were washed twice with ice-cold elution buffer (50 mM Hepes-KOH, pH 7.6, 250 mM KCl, 1 mM MgCl_2_, 1 mM EGTA, 5% glycerol, and 0.1% Tween-20). To elute the column, 250 μl of elution buffer containing 0.5 mg/ml HA dipeptide was added to the column and the flow-through fraction was collected. After a 30-min incubation, another aliquot was added. This was repeated for a total of six fractions. Elution fractions two through five were aliquoted in 10 μl vol and flash frozen on liquid nitrogen.

For the purification of PP2A^Rts1-3xHA^ (DK660) and mss4-3xHA (DK2326), a similar protocol was followed, except that cells were grown in YPD and 12 g of powder was resuspended in 30 ml of extract buffer. Mss4-3xHA cell powder was resuspended in 30 ml of extract buffer (50 mM Tris-HCl, 7.5, 700 mM NaCl, 150 mM NaF, 150 mM **β**- glycerophosphate, 1 mM EGTA, 5% Glycerol, 0.25% Tween-20, 2 mM PMSF).

### Baculovirus expression and Yck1 purification

To purify Yck1 protein from insect cells, the *YCK1* ORF was amplified (oligos: CCGGGATCCATGTCCATGCCCATAGCA and CGGCTCGAGTTAGTGGTGGTGGTGGTGGTGGTGGTGGCAACAACCTAATTTTTGGA), digested with BamHI and XhoI and ligated into a modified pFastBac-GST-TEV-MCS- 8xHIS vector (renamed to pAJ10) provided by Seth Rubin (UCSC). The resulting plasmid pAJ12 (pFastBac-GST-TEV-YCK1-8xHIS) was verified by sequencing. pAJ12 was then transformed into competent DH10 *E. coli* cells according to the product protocol to assemble the *YCK1* construct into a bacmid. Bacmid DNA was then isolated and transfected into SF9 insect cells using Fugene transfection reagent. The baculovirus was isolated from the initial transfection and different amounts of the viral titer were used to infect SF9 cells inoculated in 400 mL culture. On day 3 post-infection, cells were pelleted at 4,000 rpm for 15 min at 4°C. The pellet was resuspended in 20 mL lysis buffer (25 mM Tris, pH 8.0, 300 mM NaCl, 1 mM DTT, 5% w/v glycerol, 1 mM PMSF, 1x LPC Protease inhibitor cocktail) in 50 mL falcon tubes and kept on ice for 15 min. The suspension was then sonicated in 4x30s cycles, incubating the tubes on ice for 15 s after each sonication. The remaining extract was centrifuged at 16,000 g for 35 min at 4°C. The protein extract was then loaded onto a 5 mL glutathione-agarose column (washed and equilibrated with 15 mL lysis buffer) at ∼20 mL/h. The column was washed with 4 column volumes of wash buffer 1 (25 mM Tris, pH 8.0, 500 mM NaCl, 1 mM DTT) at ∼30 mL/h. The GST-tagged protein was eluted from the column using elution buffer 1 (40 mM Tris, pH 8.0, 200 mM NaCl, 20 mM Glutathione, 1 mM DTT). The elution buffer was loaded onto the column at ∼20 mL/h and ∼750 µL fractions were collected. The protein-containing fractions were identified by Bradford assay and the pooled protein (∼5 mL) was incubated with 50 µg TEV protease to cleave off the GST tag and incubated overnight at 4°C on a roller. On the next day, a 10 µL sample of the digested protein prep was run on a 12.5% polyacrylamide gel and stained with Coomassie to confirm complete cleavage of the GST tag. The remaining protein was loaded onto a 1.5 mL Ni-NTA^+^ column at 10 mL/h. The column was washed with wash buffer 2 (25 mM sodium phosphate buffer, pH 7.6, 500 mM NaCl, 20 mM imidazole, 5% glycerol, 0.1% Tween 20, 0.5 mM DTT) at 30 mL/h. Yck1-8xHIS was eluted using elution buffer 2 (25 mM sodium phosphate buffer, pH 8.0, 200 mM NaCl, 350 mM imidazole, 5% glycerol, 0.025% Tween 20). For elution, 250 µL elution buffer 2 was added to the column each time, the column was incubated at room temperature for 15 mins and the eluted fraction was then collected. Each fraction was tested for presence of protein using Bradford’s assay and the positive protein fractions were pooled together and dialyzed overnight at 4°C using dialysis buffer (50 mM HEPES, pH 8.0, 50 mM KCl, 1 mM MgCl_2_, 1 mM DTT, 30% glycerol).

### In vitro assays

All in vitro assays were performed with 1:10 dilutions of affinity purified protein stocks with dilution buffer (50 mM Hepes-KOH 7.6, 2 mM MgCl_2_, 0.05% Tween-20, 1 mM EGTA, 10% glycerol, 1 mM DTT, 20 μg/ml BSA). To demonstrate that PP2A^Rts1^ can oppose phosphorylation of Yck1/2 in vitro (Figure 3B), 2 μl of diluted Rts1-3xHA was incubated with 5 μl of diluted Yck1-6xHIS and 3xHA-Yck2 in the presence or absence of 2mM ATP and 50 µM okadaic acid. 30 μl reactions were carried out in assay buffer for 20 minutes at 30°C and was terminated by adding 12.5 μl of 4X protein sample buffer. To demonstrate the ability of Yck1-6xHIS to phosphorylate 3xHA-Ypk1 and Mss4-3xHA (Fig. 2B and 4D), 1 μl of diluted Yck1-6xHis was incubated with 2 μl of 3xHA-Ypk1 or 5 μl of Mss4-3xHA in a total volume of 30 μl. The reactions were incubated at 30°C for 30 min and terminated by adding 12.5 μl of 4X sample buffer. The samples were boiled at 100°C for 5 min and 20 μl of each reaction was analyzed by Western blotting.

## ACKNOWLEDGEMENTS

We thank members of the laboratory for advice and support. We also thank Ted Powers (University of California, Davis) for the Ypk-pT662 phosphospecific antibody, David Toczyski (University of California, San Francisco) for 12CA5 antibody, and Kevan Shokat (University of California, San Francisco) for providing 3-MOB PP1, 3-MB-PP1 and 3-BrB-PP1. We are grateful to Lucy Robinson (Louisiana State University Health Science Center) for strains and Seth Rubin (University of California, Santa Cruz) for reagents and helping purifying Yck1.

## COMPETING INTERESTS

The authors declare no competing or financial interests.

## FUNDING

This work was supported by National Institutes of Health grant GM131826 to DK; PAIDI 2020 (P18-FRJ-1132) co-financed by Programa Operativo FEDER 2014-2020 from Junta de Andalucía to R.L. and PAIDI 2020 (PROYEXCEL_00174) from Junta de Andalucía to MAG.

## AUTHOR CONTRIBUTIONS

Conceptualization: R.L., A.J., D.K, M.A-G; Methodology: R.L., A.J., S.A., M.A-G; Formal analysis: R.L., A.J., S.A.,D.K., M.A-G.; Writing - original draft: R.L., D.K., M.A-G.; Writing - review & editing: R.L., D.K., M.A-G; Supervision: R.L., D.K., M.A-G; Funding acquisition: R.L., D.K., M.A-G.

## DATA AVAILABILITY

Strains and plasmids are available upon request. The authors affirm that all data necessary for confirming the conclusions of the article are present within the article, figures, and tables.

---

**Supplementary Table 1.**
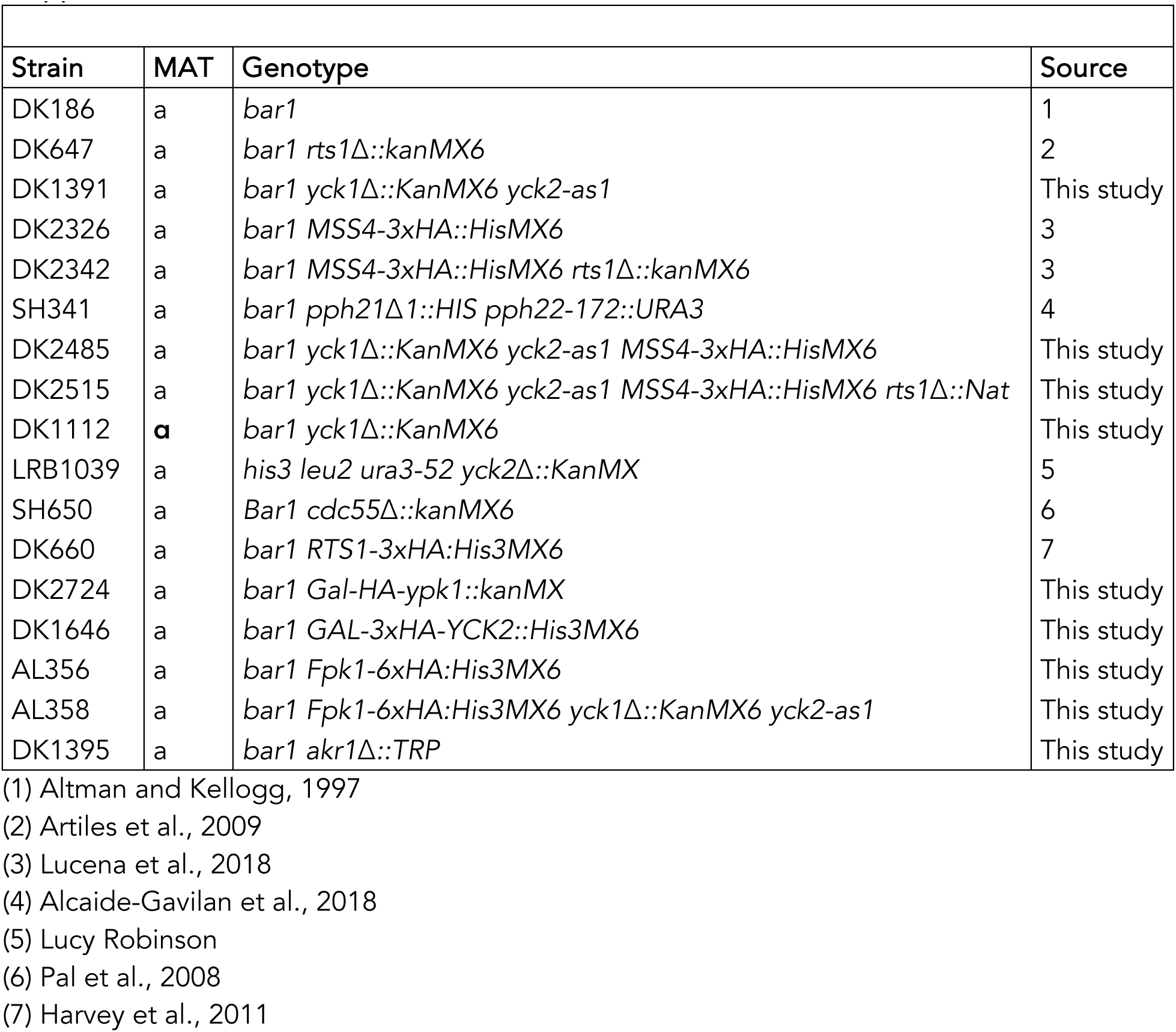
Strains used in this study.

## Notes

### Competing Interest Statement

The authors have declared no competing interest.

